# Honey bees are resilient to the long-term presence of alcohol in their diet

**DOI:** 10.1101/2024.10.31.621217

**Authors:** Monika Ostap-Chec, Daniel Bajorek, Weronika Antoł, Daniel Stec, Krzysztof Miler

**Author notes:** Correspondence: Monika Ostap-Chec, Institute of Environmental Sciences, Faculty of Biology, Jagiellonian University, Gronostajowa 7 St., 30-387 Kraków, Poland., Krzysztof Miler, Institute of Systematics and Evolution of Animals of the Polish Academy of Sciences, Sławkowska 17 St., 31-016 Kraków, Poland.

## Abstract

Previous studies on various organisms have suggested that low doses of ethanol can have stimulatory effects, while higher doses may lead to toxicity, a response known as hormesis. Low ethanol concentrations occur naturally in the environment, particularly in fermenting fruits and flower nectar, where pollinators such as honey bees may encounter it. This study aimed to investigate the potential hormetic effects of low-level ethanol consumption on honey bees. Bees were divided into three groups: one provided with only sucrose solution, one both with sucrose and 0.5% ethanol in sucrose, and one with only 1% ethanol in sucrose. The bees were exposed to these diets for 14 days, and their performance was assessed through survivorship, flight endurance, body mass, lipid content, and trehalose and ethanol levels in the haemolymph. The results showed no significant differences in most parameters between the groups. However, bees constantly exposed to 1% ethanol had slightly higher trehalose levels compared to the control group, suggesting a possible adaptive response to ethanol exposure. Ethanol levels in the haemolymph differed significantly between groups, with bees exposed to ethanol showing its detectable levels in their system. While no clear hormetic effects were observed in terms of improved performance, the elevated trehalose levels in bees constantly exposed to 1% ethanol may indicate adaptations protecting from ethanol-induced damage. The study provides insights into how honey bees tolerate low-level ethanol exposure and highlights the need for further research on the ecological implications of ethanol consumption in pollinators.

## Introduction

Hormesis is a dose-response phenomenon in which low levels of environmental agents stimulate beneficial effects, while higher doses become inhibitory or toxic (Stebbing 1982, Mattson 2008). This concept applies across various taxa and factors, such as temperature, water availability, and nutrition (Parsons 2001). Hormesis is considered a general phenomenon resulting from evolutionary adaptations to the levels of environmental factors present in the habitat (Parsons 2001). At the cellular level, hormetic responses are driven by signalling pathways and molecular mechanisms that activate adaptive stress responses (Mattson 2008, Mattson and Calabrese 2010).

In agroecosystems, hormetic effects are observed in insects exposed to sub-lethal doses of insecticides, leading to enhanced reproduction, survival, longevity, growth, and stress tolerance (Cutler et al. 2022). For example, green peach aphids exposed to low concentrations of imidacloprid exhibited increased reproduction and population growth (Ayyanath et al. 2013). Similar responses have been reported in other insect pests (Haddi et al. 2016, Rakotondravelo et al. 2019). These hormetic responses involve molecular and biochemical mechanisms, including increased expression of heat shock proteins (HSP), detoxification genes, and changes in antioxidant activity (Rix and Cutler 2022). The implications of hormesis extend beyond pest management to pollinator conservation, as stimulatory effects have also been observed in bees exposed to low doses of pesticides (Cutler and Rix 2015). However, further research is needed to understand the ecological consequences of hormesis in agroecosystems (Cutler et al. 2022).

Types of factors and environments that promote hormetic effects also need further study. Ethanol is relevant in this context as it occurs naturally in flower nectar and fruits due to yeast fermentation of sugars (Dudley and Maro 2021). This process dates back to the Cretaceous period (Dudley 2014), hence many insects that rely on such fermenting resources may have evolved adaptive responses to ethanol exposure. For instance, fruit flies, which use ethanol-spiked substrates for feeding and egg-laying, exhibit hormesis in response to ethanol itself and acetaldehyde, a metabolic byproduct of ethanol (Starmer et al. 1977, Parsons et al. 1979, Parsons 1989). Such adaptations illustrate how certain species thrive in ethanol-containing environments (Devineni and Heberlein 2013). Ethanol exposure seems particularly relevant in agroecosystems such as orchards, where ripening fruits can contain ethanol at varying concentrations. Some yeast strains isolated from orchard soils have been shown to tolerate ethanol concentrations as high as 20% (Moneke et al. 2008), suggesting that high ethanol levels may occur in these habitats.

Pollinators such as honey bees also highlight potential dose-dependent responses to ethanol. For example, some studies on bees (Abramson et al. 2000, Black et al. 2021) show that high doses of ethanol – 5% and above – lead to greater behavioural impairment in learning performance than lower doses (1 or 2.5%). Hranitz et al. (2010) reported that HSP levels in the brain tissues of bees increased with feeding on ethanol up to 5%, but the adaptive stress response was inhibited at 10% ethanol, which suggested that beyond certain levels bees were no longer able to cope with ethanol exposure. Mustard et al. (2019) also found that bees preferred to consume solutions with only relatively low ethanol concentrations of 1.25% to 2.5% and that higher levels increased bees’ mortality. These findings show that while bees tolerate low ethanol exposure, higher doses impair performance and survival. Similarly to fruit flies, honey bees may encounter ethanol in their environment as they utilise flower nectar and ripe fruit juices. It seems reasonable to expect they show hormetic patterns in response to ethanol.

However, research on the effects of low-level ethanol exposure in bees remains underrepresented, making it difficult to establish whether there are any beneficial effects under such conditions. Ahmed et al. (2022) suggest that hormetic effects may be possible across a range of ethanol concentrations (1%, 2.5%, and 5%) in terms of bees’ flight kinematics. At 1% ethanol, unlike at 2.5% and 5%, flight parameters were not impaired in bees but, at the same time, were qualitatively different from the control. Ostap-Chec et al. (2024) further suggest that the frequency of ethanol encounters plays a significant role in bees’ mortality, with more frequent consumption of 1% ethanol leading to higher mortality over time. Therefore, whether ethanol feeding benefits bees in any way remains an open question. A key gap in investigating this issue lies in determining the ethanol frequencies and concentrations that align with an eco-evolutionary hormetic scenario in bees and other pollinators.

In this study, we explore the potential beneficial effects of low-level ethanol consumption in bees. We exposed bees to a two-week dietary regime, comparing their performance across three groups: bees with no ethanol exposure, bees that self-dosed with 0.5% ethanol, and bees that were forced to constantly consume 1% ethanol. We hypothesised that bees allowed to self-dose with low ethanol concentrations would outperform the other groups. To assess performance we measured survivorship, flight endurance, body and lipid mass, and sugar reserves in the haemolymph. We also measured ethanol levels in the haemolymph to confirm that the diet differentiated bees in terms of ethanol presence in their systems.

## Materials and Methods

### General procedure

The experiment was performed in four replicates, with a two-day delay between the preparation of each replicate. For each replicate, newly emerged bees (*Apis mellifera carnica* Poll.) were obtained from two unrelated, queen-right colonies with naturally inseminated queens. For that purpose, we took a bee-free frame with capped brood from each colony and placed it in an incubator (KB53, Binder, Germany) at 32 °C overnight. The next morning, all emerged bees were marked with a coloured dot on the thorax using a non-toxic paint marker, and released into an unrelated hive. The hive environment supports proper development and strengthens the bees’ immune system, which results in high subsequent survival rates in laboratory cages, allowing for extended experiments (Ostap-Chec et al. 2021, Ostap-Chec et al. 2024). At seven days old, the marked bees were recollected from the hive using forceps, placed in wooden cages, and transported to the laboratory for a three-day acclimation period. This acclimation ensured the bees remained naïve to ethanol in their immediate diet before the experiment. Then, at ten days old, the experimental period began, involving adult bees close to the age when they typically initiate foraging (i.e., around day 12 of life) (Dukas 2008).

In the laboratory, bees were kept in wooden cages, with 50 individuals per cage, in an incubator (KB400, Binder, Germany) set at 32 °C. Bees were given *ad libitum* access to water and food. During the acclimation period, all cages were provided with 1 M (i.e. 22% w/w) sucrose solution, while the diet varied during the experimental period depending on the group (see below). To avoid evaporation and fermentation, fresh food was provided twice daily, around 8 AM and 8 PM. Water was also refreshed daily, and any dead bees were noted and removed. In each replicate, 12 cages were prepared: six for flight performance and body mass assessment, and six for trehalose level analysis and sobriety tests. The number of live bees in each cage was recorded at the start of the experimental period to track survivorship until the study’s termination.

### Flight performance and body mass

The six cages in each replicate designated for flight performance and body mass assessment were divided into three groups, each with two cages. The groups differed in their diet during the experimental period. In group 1 (G1), bees received two feeders with only 1 M sucrose solution. In group 2 (G2), bees received 1 M sucrose solution (one feeder) and a sucrose solution containing 0.5% ethanol (another feeder). In group 3 (G3), bees received two feeders with only a sucrose solution containing 1% ethanol. The ethanol-supplemented solutions were prepared to be iso-caloric with the 1 M sucrose solution by reducing the sucrose content by 1.38 g for each mL of ethanol (assuming 5.5 kcal per 1 mL of ethanol) to prevent confounding effects related to ethanol’s caloric value when digested (Miler et al. 2021a). The bees were exposed to these diets for 14 days, after which flight performance and body mass were assessed.

One cage from each group was assigned for flight performance assessment using flight mills (Biospekt, Poland) (Brodschneider et al. 2009). Each bee designated for flight test had an opalith plaque glued to its thorax the day before testing, using shellac, to allow tethering to the flight mill the following day. Bees were then tested individually on the flight mill (Blanken et al. 2015), with one bee per group tested simultaneously, and each test began immediately after tethering. If a bee did not initiate flight within one minute, the test was terminated. If a flight was initiated, the test continued until the first stop. The flight speed was recorded automatically on a connected laptop every two seconds. In total, we tested 82, 69, and 67 bees from G1, G2, and G3, respectively, across all replicates.

The remaining cage from each group was used for body mass analysis. Bees were frozen, transferred to labelled Eppendorf tubes and dried at 60 °C for 96 hours in a laboratory drier (ED56, Binder, Germany). Each bee was weighted to obtain its dry mass using a laboratory balance. Bees were then immersed in diethyl ether for 48 hours to dissolve and extract lipids, followed by a fresh ether wash and an additional drying period of 96 hours at 60 °C. The lipid-free mass was then recorded. Lipid content was calculated by subtracting the lipid-free mass from the dry mass (as in Colinet et al. 2006, Wilson-Rich et al. 2008, Visser and Ellers 2012). Dry mass was obtained from 93, 104, and 105 bees from G1, G2, and G3, respectively (all replicates). Due to material fragility, reliable lipid-free mass measurements were obtained for 46, 42, and 54 bees from G1, G2, and G3, respectively (all replicates).

### Trehalose level analysis and sobriety tests

The same procedure was followed for another set of cages. After 14 days of dietary exposure, haemolymph was sampled from the bees on the final day of the experiment using the antennae method (Borsuk et al. 2017). Each bee was placed on a Styrofoam plate, and one antenna was cut with micro scissors. Gentle pressure was applied to the abdomen to collect haemolymph in an end-to-end 10 μl microcapillary tube, which was then stored in a cryotube at −20 °C. Haemolymph from one cage per group was used for trehalose level analysis, while another cage provided haemolymph for sobriety tests.

Trehalose levels were measured using a trehalose assay kit (K-TREH, Neogen, USA), following the manufacturer’s protocol. A minimum of 1 μl of each haemolymph sample was diluted 200x with distilled water. If the individual sample volume was insufficient, samples were paired within the same experimental group and colony. Samples that failed to achieve the minimal volume even after pairing were discarded as volumes below 1 µl were considered not precise enough to be used in quantitative analysis. For the analysis, 20 µl of each diluted sample was used and run in duplicate. Analysis was performed in batches (consecutive plates), always with a trehalose standard dilution series covering the range of 0.00625 – 0.4 g/L. Overall 55, 35, and 42 samples from G1, G2, and G3, respectively (all in duplicates) were analysed.

Ethanol levels were measured using an ethanol assay kit (K-ETOH, Neogen, USA), following the manufacturer’s protocol. 3 µl of haemolymph per sample was used in the analysis (the minimal volume observed in previous tests to result in detectable levels of haemolymph ethanol), filled up to 10 µl (sample volume required in the reaction) with distilled water. Most of the samples were pooled in pairs to achieve the minimal volume. Each analysis batch (plate) included an ethanol standard dilution series covering the range of 0.0125 – 0.1 g/L. Overall 32, 36, and 26 samples from G1, G2, and G3, respectively (all in duplicates) were analysed.

Absorbance increase caused by enzymatic reaction products was measured at 340 nm using the Multiskan FC microplate reader (Thermo Scientific, USA). Trehalose and ethanol levels were calculated based on respective calibration curves and recalculated to original concentrations, based on the sample dilution factor.

### Statistics

All statistical analyses were performed using R (R Core Team 2014), with plots generated using the ‘ggplot2’ package (Wickham 2016).

For survivorship analysis, we used a Cox mixed-effects model (‘coxme’ and ‘survival’ packages, Therneau 2022, 2023). The initial number of bees in each cage at the start of the diet was used as the starting point, with mortality tracked until day 13 (day 14 was excluded as in some cages bees were manipulated on day 13 in preparation for flight performance tests). Bees alive on the last day were censored. The group was included as a fixed factor, with the cage nested in the replicate as random factors. The proportional hazards assumption was tested using the cox.zph function and met.

We used a generalized linear mixed-effects model (GLMM, binomial with logit link) fitted using the ‘glmmTMB’ package (Brooks et al. 2017) to analyse flight initiation in flight performance tests. Bees that did not fly were assigned “0”, and those that flew were assigned “1”. To analyse flight parameters, GLMMs were also run, but with the following error structures: negative binomial and log link for total flight time, Gaussian with inverse link for average speed, and Gamma with log link for distance travelled. In these three models, group-specific dispersion (dispformula function) was included to account for overdispersion in the response variables. For the analysis of mass, data was sqrt-transformed and GLMMs with Gaussian error structures and identity links were run for the dry mass and mass of lipids. These two models allowed for a group-specific dispersion like those run to analyse flight parameters. All GLMMs included the group as a fixed factor and the replicate as a random factor.

To analyse trehalose levels, a linear mixed-effects model (Gaussian with identity link) was fitted using the ‘nlme’ package (Pinheiro et al. 2024). The model included the group as a fixed factor and the replicate as a random factor. As the material was divided into several analysis batches, we also included another random factor, the plate. We allowed the residual variance to differ between groups using the varIdent function. Ethanol levels in haemolymph were analysed using a zero-inflated generalized mixed-effects model fitted with the ‘glmmTMB’ package (Brooks et al. 2017), using a Gamma error structure (log link). The group was included as a fixed factor, and the replicate and plate as random factors.

The significance of the group effect was tested using the Anova function (‘car’ package, Fox and Weisberg 2019). In addition, marginal R^2^ was calculated to estimate the variance explained by the group using the ‘MuMIn’ and ‘performance’ packages (Lüdecke et al. 2021, Bartoń 2024). Models were diagnosed for proper performance and fit with the ‘DHARMa’ package (Hartig et al. 2024).

## Results

We found no significant differences in survivorship between the groups. On the last day, the survival probability (95% CI) was at 65% (61-68%) for G1, 59% (55-62%) for G2, and 60% (57-63%) for G3. Hazard ratios for G2 and G3 were not significantly different from the ratio observed in G1 (z = 1.140, p = 0.250 and z = 0.850, p = 0.400, respectively).

The probability of initiating flight was similar across the groups (χ^2^ = 1.072, df = 2, p = 0.585). Furthermore, among the bees that initiated flight, no differences were observed in total flight time (χ^2^ = 0.627, df = 2, p = 0.731, Fig. 1), average speed (χ^2^ = 0.369, df = 2, p = 0.832, Fig. 1), or distance travelled (χ^2^ = 0.840, df = 2, p = 0.657, Fig. 1).

**Fig. 1.**
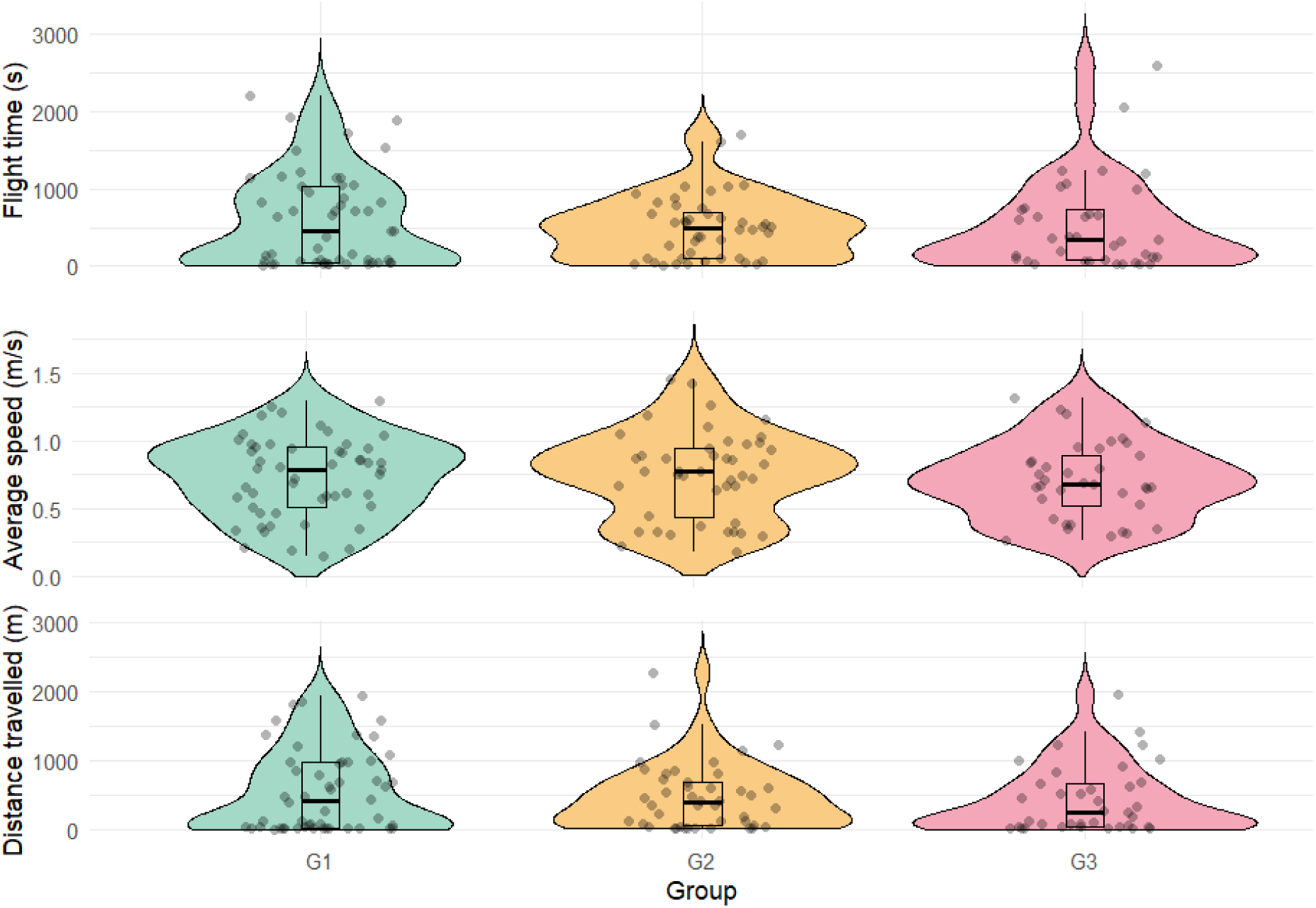
The total flight time (top panel), average speed (middle panel), and distance travelled (bottom panel) during the flight performance tests in bees without access to ethanol (G1), those with voluntary consumption of 0.5% ethanol (G2), and those forced to consume 1% ethanol (G3). Diets respective for each group lasted for two weeks before tests. Grey dots indicate individual data points. Boxplots show medians (lines), interquartile range (boxes), and the smallest and largest values within 1.5 of the interquartile range (whiskers). Coloured shadings show the kernel density estimations trimmed at 0.

We found no differences in the dry body mass (χ^2^ = 1.456, df = 2, p = 0.483, Fig. 2) or in lipid mass (χ^2^ = 0.609, df = 2, p = 0.738, Fig. 2) between the groups.

**Fig. 2.**
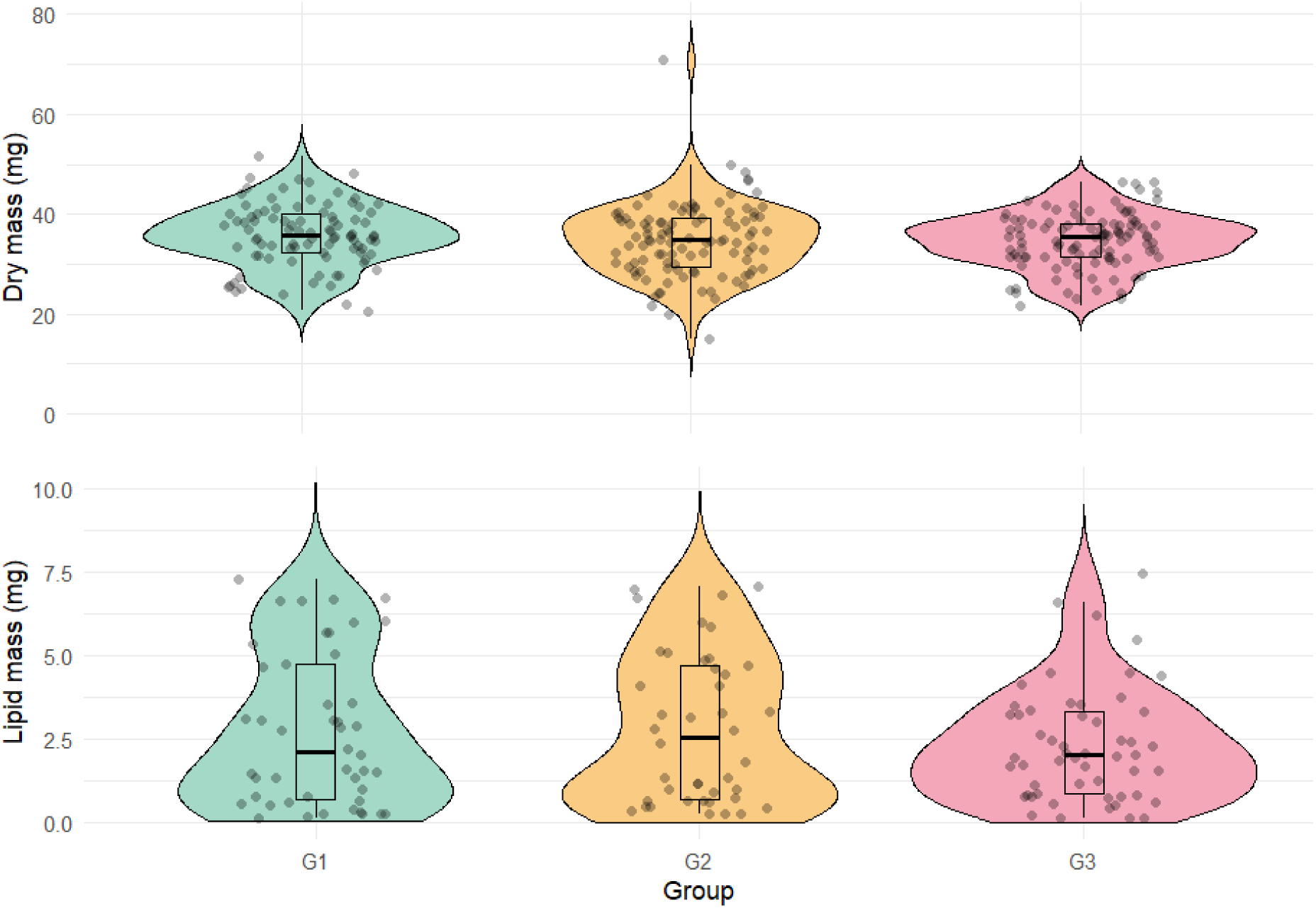
Dry mass (top panel) and lipid mass (bottom panel) of bees without access to ethanol (G1), those with voluntary consumption of 0.5% ethanol (G2), and those forced to consume 1% ethanol (G3). Diets respective for each group lasted for two weeks before mass measurements. Grey dots indicate individual data points. Boxplots show medians (lines), interquartile range (boxes), and the smallest and largest values within 1.5 of the interquartile range (whiskers). Coloured shadings show the kernel density estimations trimmed at 0.

The mean trehalose level in the control group was 22.37 ± 3.00 g/L (estimate ± SE). There was a trend toward differences in trehalose levels among the groups (F_2,106_ = 2.807, p = 0.065). Specifically, bees in G3, which had constant access to ethanol, showed higher haemolymph trehalose levels compared to G1 (estimate ± SE = 5.99 ± 2.81, t = 2.129, p = 0.036, Fig. 3). No significant difference in trehalose levels was observed between G1 and G2 (estimate ± SE = 4.85 ± 3.04, t = 1.598, p = 0.113, Fig. 3). The group explained 4.9% of the variance in trehalose levels.

**Fig. 3.**
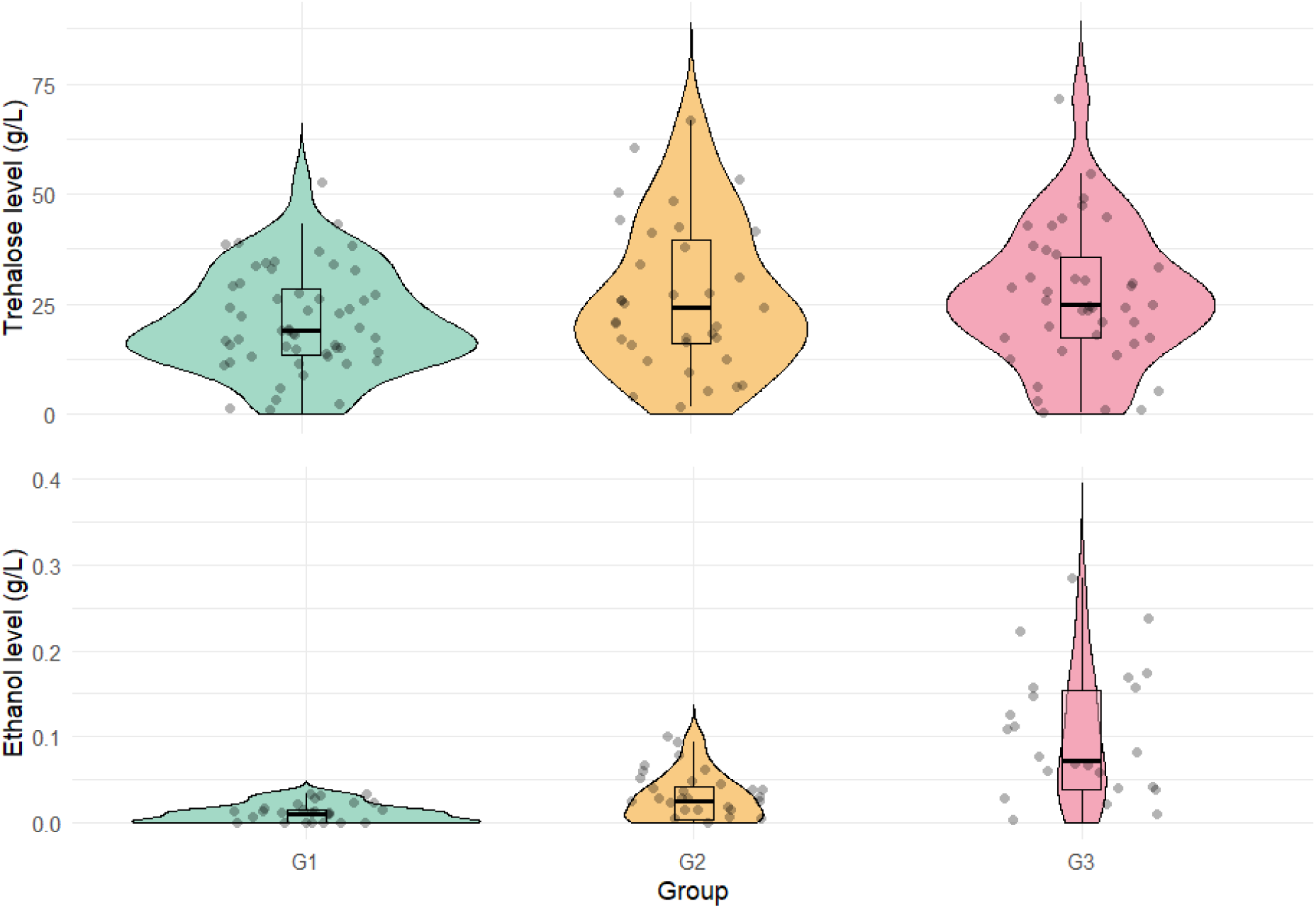
Trehalose (top panel) and ethanol (bottom panel) levels (g/L) in the haemolymph of bees without access to ethanol (G1), those with voluntary consumption of 0.5% ethanol (G2), and those forced to consume 1% ethanol (G3). Diets respective for each group lasted for two weeks before haemolymph sampling. Grey dots indicate individual data points. Boxplots show medians (lines), interquartile range (boxes), and the smallest and largest values within 1.5 of the interquartile range (whiskers). Coloured shadings show the kernel density estimations trimmed at 0.

Ethanol levels in the haemolymph differed significantly between the groups (χ^2^ = 59.383, df = 2, p < 0.001). Bees in G3, with constant ethanol access, had fewer cases of undetectable ethanol levels compared to G1 (estimate ± SE = −1.974 ± 0.822, z = −2.403, p = 0.016, Fig. 3). This pattern was not observed in G2, where bees voluntarily consumed low levels of ethanol (estimate ± SE = −0.588 ± 0.531, z = −1.108, p = 0.268, Fig. 3). In terms of non-zero values of ethanol, both G2 (estimate ± SE = 0.873 ± 0.218, z = 4.000, p < 0.001, Fig. 3) and G3 (estimate ± SE = 1.745 ± 0.227, z = 7.705, p < 0.001, Fig. 3) had significantly higher ethanol content than G1. The group explained 43.3% of the variance in ethanol levels.

## Discussion

In this study, we did not observe the expected hormetic differences between bees with no ethanol exposure, those that self-dosed with 0.5% ethanol, and those forced to constantly consume 1% ethanol. Across survivorship, flight endurance, body mass, and lipid content, we found no significant differences between the three groups. The only difference was in the haemolymph sugar reserves, where bees constantly exposed to 1% ethanol had slightly higher trehalose levels compared to the controls. Trehalose, the primary sugar in insect haemolymph, plays a crucial role in energy balance (Mayack and Naug 2010). Its level in the honey bee haemolymph is less stable compared to other sugars and depends on e.g. metabolic rate (Blatt and Roces 2001) and time after the last feeding (Mayack and Naug 2010), especially in forager-age individuals (Mayack et al. 2019), which could contribute to the high variance observed here within groups.

Apart from being an energy reserve, trehalose has other important functions which could explain its slightly elevated concentration in the group forced to constantly consume 1% ethanol. For example, in ethanol-rich environments like those inhabited by yeasts, trehalose accumulation helps protect cells from ethanol stress. In *Saccharomyces cerevisiae*, trehalose correlates with increased ethanol tolerance by safeguarding endocytosis from ethanol inhibition and protecting cellular proteins and membranes from ethanol-induced damage (Lucero et al. 2000, Bandara et al. 2009). Trehalose plays similar protective roles in other yeast species (Li et al. 2023) and insects, where it is involved in stress protection and recovery (Shukla et al. 2015). Genetic studies in *Drosophila* have shown that trehalose-related genes are critical for normal development, body water homeostasis, and desiccation tolerance (Yoshida et al. 2016). Trehalose metabolism is regulated by complex mechanisms, including transcription factors, enzyme modifications, and neuroendocrine control (Tellis et al. 2023). Understanding its role in insects can offer potential applications in pest control strategies and biotechnology (Shukla et al. 2015, Tellis et al. 2023).

Ethanol often induces hormetic effects, a response seen across various organisms (Calabrese and Baldwin 2003). Although we did not observe such effects in this study, our results suggest that honey bees are resilient to ethanol in their diet. Remarkably, two weeks of continuous ethanol consumption had virtually no impact on survivorship, flight endurance, body mass, or lipid reserves. This finding contrasts with earlier studies (Rassmussen et al. 2021, Ostap-Chec et al. 2024) that showed increased mortality with 1% ethanol in the diet, but these studies used much older bees than we did. The bees in those studies were either adult winter bees of mixed ages (Rassmussen et al. 2021) or almost two weeks older than in our study (Ostap-Chec et al. 2024). This suggests that the toxic effects of dietary ethanol may be more pronounced for older bees. However, our study reflects the typical lifespan of worker bees, which begin foraging around two weeks old and generally live about another two weeks (Dukas 2008). Therefore, our results are highly relevant to the life cycle of most worker bees.

The detected resilience to ethanol suggests that honey bees may have evolved adaptations to ethanol encounters, such as those with fermenting flower nectar or over-ripe fruit juices. The presence of ethanol in wild fruits, typically up to around 1%, has been documented (Eriksson and Nummi 1982, Dudley 2002, 2004, Amato et al. 2020, Campbell et al. 2022, Casorso et al. 2023). Ethanol is also thought to be routinely present in flower nectar (Lievens et al. 2015), with studies confirming its presence in this type of resource (Kevan et al. 1988, Ehlers and Olesen 1997, Jakubska et al. 2005, Goodrich et al. 2006, Wiens et al. 2008, Golonka et al. 2014, Gochman et al. 2016, Rering et al. 2018). However, much of this research is anecdotal, and further studies on the concentrations and frequency of ethanol encounters in honey bees and other pollinators are needed. Determining this is necessary to investigate ethanol hormesis in these insects and design realistic exposure scenarios in the laboratory. It remains an open question whether human-altered environments, such as orchards or vineyards, create a mismatch with evolutionarily typical ethanol exposures and responses in insects.

Fruit flies are among the most well-known insects that utilise ethanol-rich habitats, and striking parallels can be drawn between fruit flies and honey bees. In *Drosophila melanogaster* larvae, ethanol exposure induces a significant increase in alcohol dehydrogenase (ADH) activity and ADH mRNA transcripts, which allows larvae to use ethanol as a resource (Geer et al. 1988). However, this process does not occur in adult flies (Geer et al. 1988). While the effects of ethanol on honey bee larvae have not been studied, adult honey bees, like adult fruit flies, do not show signs of ADH induction under ethanol exposure (Miler et al. 2022, Ostap-Chec et al. 2024). Fruit flies have emerged as invertebrate models for studying the genetic and physiological mechanisms underlying alcohol-related behaviours, offering insights applicable to human alcoholism research (Scholz and Mustard 2013). They demonstrate tolerance and dependence in response to chronic ethanol exposure (Berger et al. 2004, Devineni and Heberlein 2009, Lin and Rankin 2012), and similar reactions are observed in honey bees (Miler et al. 2018, Ostap-Chec et al. 2021). Both taxa show behavioural responses to ethanol that mirror those seen in mammals, such as altered locomotion (Heberlein et al. 2004, Miler et al. 2018b, Nuñez et al. 2023). However, many other insects, such as butterflies, ambrosia beetles, hornets and bumblebees, also use ethanol-spiked habitats (Dierks and Fischer 2008, Ranger et al. 2018, Jones and Agrawal 2022, Bouchebti et al. 2024) and may offer interesting comparative perspectives for future studies.

There is growing evidence that hormesis occurs across a wide range of organisms, from bacteria to vertebrates, in response to numerous chemical and environmental stressors (Constantini et al. 2010, Berry and López-Martínez 2020). As we show here, the concept is highly relevant to ecological and evolutionary processes. We investigated the potential hermetic effects of low-level ethanol exposure in honey bees, comparing bees that self-dosed or were forced to consume ethanol with controls. Although we found no significant differences in survivorship, flight endurance, or body and lipid mass across the groups, bees exposed to constant ethanol showed slightly elevated trehalose levels, suggesting a possible protective mechanism against ethanol-induced stress. The results indicate that honey bees are resilient to low-level ethanol exposure, though further research is needed to fully explore the ecological and physiological implications of this finding.

## Acknowledgements

This work was supported by the National Science Centre, Poland [grant number Sonata UMO-2021/43/D/NZ8/01044].

## Author Contribution Statement

MO-C: Conceptualization; Investigation; Methodology; Writing—original draft; Writing— review & editing. DB: Investigation; Methodology; Software; Writing—review & editing. WA: Formal analysis; Investigation; Methodology; Validation; Writing—review & editing. DS: Resources; Writing—review & editing. KM: Conceptualization; Data curation; Formal analysis; Funding acquisition; Investigation; Methodology; Project administration; Resources; Software; Supervision; Visualization; Writing—original draft; Writing—review & editing.

